# The proxiomes of CDJ5 and PGRL1 in *Chlamydomonas reinhardtii* overlap depending on the ability of CDJ5 to bind a 4Fe-4S cluster

**DOI:** 10.1101/2025.08.22.671721

**Authors:** Katharina König, Melissa Misir, Justus Niemeyer, Sarah Gabelmann, Saskia Zeilfelder, Frederik Sommer, Michael Schroda

## Abstract

The chloroplast chaperone HSP70B from *Chlamydomonas reinhardtii* works with the J-domain co-chaperones CDJ1 to CDJ6. CDJ1 delivers unfolded proteins, while CDJ2 delivers VIPP1 as substrate to HSP70B. CDJ3 to CDJ5 contain 4Fe-4S clusters in addition to the J domain, but their function is unknown. To investigate the function of CDJ5, we performed TurboID - mediated proximity labeling on wild-type CDJ5 (CDJ5-WT) and CDJ5 mutants with an impaired ability to stimulate HSP70B’s ATPase activity (CDJ5-AAA) or bind a 4Fe-4S cluster (CDJ5- SSS). Our results revealed that the proxiomes of all CDJ5 variants contained HSP70B and HSP90C. Furthermore, the proxiomes of CDJ5-WT and CDJ5-AAA overlapped extensively but differed from that of CDJ5-SSS, suggesting that the localization of CDJ5 to a chloroplast microcompartment depends on the presence of a functional 4Fe-4S cluster or its redox state. The CDJ5-WT and CDJ5-AAA proxiomes were enriched with proteins that regulate photosynthetic electron flow or are involved in the biogenesis of thylakoid membrane protein complexes and pigments. These proteins were also present in the proxiome of PGRL1 found in the CDJ5-WT and CDJ5-AAA proxiomes. Overall, our results suggest that CDJ5 acts with HSP70B/HSP90C via its 4Fe-4S cluster to regulate photosynthetic electron flow and thylakoid membrane protein complex biogenesis.

**Highlight:** Depending on its 4Fe-4S cluster, HSP70B co-chaperone CDJ5 localizes to a chloroplast microcompartment defined by PGRL1 where it could act in regulating photosynthetic electron flow and thylakoid membrane protein complex biogenesis.

## Introduction

Molecular chaperones of the heat shock protein (HSP) 70 class use the energy of ATP hydrolysis to induce conformational changes in substrate proteins. This is key to the function of HSP70s in assisting the folding of unfolded proteins to the native state, in driving the translocation of proteins across membranes, or in facilitating the assembly and disassembly of protein complexes (Rosenzweig *et al*., 2019). Essential for mediating substrate specificity of HSP70s are the J-domain proteins (Craig *et al*., 2006). These bind specific substrates via specialized domains that they harbor in addition to the J domain and deliver these substrates to the substrate-binding domain of their HSP70 partner while concomitantly triggering ATP hydrolysis. The stimulation of ATP hydrolysis depends on the highly conserved histidine-proline-aspartate (HPD) tripeptide located in the loop between helix II and III of the J domain (Tsai and Douglas, 1996; Wall *et al*., 1994). Thus, specific functions of an HSP70 in a cellular compartment are mediated by its set of J-domain proteins. Of the ∼63 J-domain proteins encoded by *Chlamydomonas*, only six (chloroplast DnaJ homologues CDJ1-6) are considered to be targeted to the chloroplast where they determine the functional specificity of chloroplast HSP70B (Schroda and deVitry, 2023). CDJ1 delivers unfolded protein substrates for folding assistance by HSP70B (Veyel *et al*., 2014; Willmund *et al*., 2008). CDJ2 delivers the VIPP1 protein to HSP70B for driving the (dis)assembly of VIPP1 oligomers (Liu *et al*., 2007; Liu *et al*., 2005). CDJ6 contains a tetratricopeptide domain in addition to its J domain but its function is unknown. CDJ3-5 are very weakly expressed proteins that contain bacterial-type ferredoxin domains with 4Fe-4S clusters in addition to their J domains (Dorn *et al*., 2010). Proteins with this domain architecture exist in chloroplasts as well as in mesophilic archaea (Thaumarchaeota), where they were introduced by horizontal gene transfer 750-900 million years ago (Petitjean *et al*., 2012). Phylogenetically, CDJ3 and CDJ4 belong to a different clade than CDJ5. CDJ3/4 were shown to interact with HSP70B in the ATP state and to stimulate ATP hydrolysis of HSP70B but unlike CDJ1 failed to assist in the refolding of unfolded proteins (Dorn *et al*., 2010; Veyel *et al*., 2014). CDJ3 and CDJ4 were demonstrated to contain redox-active 4Fe-4S clusters and CDJ3 was found in stromal complexes of 550 to 2800 kDa that contain RNA (Auerbach *et al*., 2017; Dorn *et al*., 2010). Virtually nothing is known about CDJ5. However, of the eight *cdj5* mutant strains available in the *Chlamydomonas* library project (CLiP) (Li *et al*., 2016), seven contain an insertion of the mutagenesis cassette in the 3’UTR and one in an intron (https://www.chlamylibrary.org/), suggesting that *CDJ5* could be an essential gene.

To shed light onto the function of CDJ5, we applied our previously established TurboID-based proximity-labeling tool (Kreis *et al*., 2023a) to CDJ5 to obtain information about proteins in its proximity. We show that the CDJ5 proxiome largely overlaps with that of PGRL1, a protein involved in cyclic electron flow (CEF), and that this depends on the ability of CDJ5 to bind a 4Fe-4S cluster. Hence, our results suggest a regulatory role of CDJ5 mediated by its 4Fe-4S cluster in photosynthetic electron flow and in thylakoid membrane protein complex biogenesis.

## Materials and Methods

### Strains and cultivation conditions

*Chlamydomonas reinhardtii* UVM4 cells (Neupert *et al*., 2009) were grown mixotrophically in Tris-Acetate-Phosphate (TAP) medium (Kropat *et al*., 2011) on a rotatory shaker at 25°C and constant light intensity of ∼30-60 μmol photons m^−2^ s^−1^. For biotin treatments, exponentially growing cells (2-4 x 10^6^ cells mL^-1^) were incubated with 1 mM biotin at 25°C under constant light intensity of 30 µmol photons m^−2^ s^−1^ for 4 h. Transformation was performed via the glass beads method (Kindle, 1990) as described previously (Hammel *et al*., 2020) with constructs linearized via digestion with NotI (CDJ5-3xHA construct) or EcoRV (all other constructs). Transformants were selected on TAP medium containing 100 µg mL^-1^ spectinomycin or 10 µg mL^-1^ paromomycin. Cell densities were determined using a Z2 Coulter Counter (Beckman Coulter).

### Plasmid constructions

*Level 0 constructs –* The cDNA sequence encoding CDJ5 (Cre07.g320350) with its predicted chloroplast transit peptide was interrupted by the three *Chlamydomonas RBCS2* introns at three AG/G sites according to Baier *et al*. (2018). The sequence was split into two fragments flanked with BbsI recognition sites, synthesized by IDT as gBlocks and assembled into the pAGM1287 vector in a restriction-ligation reaction with BbsI and T4 DNA ligase, giving pMBS618. Site-directed mutagenesis on pMBS618 using primers #1780 and #1781 (Supplementary Table S1) resulted in the CDJ5-AAA variant (pMBS619), and primers #1782 and #1783 in the CDJ5-SSS variant (pMBS620). The coding sequence for LCP1 (Cre02.g141550) including its predicted chloroplast transit peptide was reverse translated using the most-preferred *Chlamydomonas* codons. The sequence was interrupted by the first two *Chlamydomonas RBCS2* introns at two AG/GC sites, split into two fragments flanked by BbsI recognition sites and synthesized by IDT as gBlocks. The two gBlocks were then assembled into the pAGM1287 vector, yielding pMBS1050. Similarly, the coding sequence for the PSBQ domain-containing protein (QCP1, Cre03.g169900) containing its predicted chloroplast transit peptide, was reverse translated using the most-preferred *Chlamydomonas* codons. The *Chlamydomonas RBCS2* intron 1 was inserted twice, and intron 2 once at three AG/GC sites. The sequence was split into two fragments with BbsI recognition sites, synthesized by IDT and assembled into the pAGM1287 vector giving pMBS1049. The construction of the level 0 part pMBS1048 for PGRL1 (Cre07.g340200) is described in Kreis *et al*. (2023a). All constructs represent level 0 parts for the B3/4 position according to the Modular Cloning (MoClo) syntax for plant genes (Patron *et al*., 2015; Weber *et al*., 2011). Correct cloning was verified by Sanger sequencing (Microsynth Seqlab GmbH).

*Level 1 and level 2 constructs –* The constructed level 0 parts were used together with level 0 parts (pCM) from the *Chlamydomonas* MoClo toolkit (Crozet *et al*., 2018) and with new level 0 parts (pMBS) to fill the respective positions in level 1 modules as follows: A1-B1 – pCM0-015 (*HSP70A-RBCS2* promoter + 5’ UTR); A1-B2 – pCM0-020 (*HSP70A-RBCS2* promoter + 5’ UTR); A1-B1 – pCM0-010 (*PSAD* promoter + 5’UTR); B2 – pCM0-052 (*PSAD chloroplast transit peptide*); B3-B4 – pMBS618 (*CDJ5-WT*), pMBS619 (*CDJ5-AAA*), pMBS620 (*CDJ5- SSS*), pMBS1050 (*LCP1*), pMBS1049 (*QCP1*), pMBS1048 (*PGRL1*) or pMBS1195 (*mNeonGreen*); B5 – pCM0-100 (*3xHA*), pMBS512 (*TurboID-C*), pMBS1076 (*mNeonGreen*); B6 – pCM0-119 (*RPL23* terminator + 3’ UTR). The respective level 0 parts for the CDJ5 constructs with 3xHA and the destination vector pICH47742 (Weber *et al*., 2011) were combined with BsaI and T4 DNA ligase and assembled into the level 1 modules CDJ5-WT- 3xHA (pMBS627), CDJ5-AAA-3xHA (pMBS628) and CDJ5-SSS-3xHA (pMBS629). The level 1 modules were then combined with pCM1-01 (level 1 module with the *aadA* gene conferring resistance to spectinomycin flanked by the *PSAD* promoter and terminator) from the *Chlamydomonas* MoClo kit, with plasmid pICH41744 containing the correct end-linker, and with destination vector pAGM4673 (Weber *et al*., 2011), digested with BbsI, and ligated with T4 DNA ligase to generate level 2 devices CDJ5-WT-3xHA (pMBS636), CDJ5-AAA-3xHA (pMBS637) and CDJ5-SSS-3xHA (pMBS638). For CDJ5 fused with TurboID or mNeonGreen (CDJ5-WT_TID_, pMBS1036; CDJ5-AAA_TID_, pMBS1037; CDJ5-SSS_TID_, pMBS1038; CDJ5-WT_mNG_, pMBS1209), LCP1 with TurboID or mNeonGreen (LCP1_TID_, pMBS1042; LCP1_mNG_ pMBS1241), QCP1 with TurboID or mNeonGreen (QCP1_TID_, pMBS1040; QCP1_mNG_ pMBS1240), PGRL1 with TurboID or mNeonGreen (PGRL1_TID_, pMBS1045; PGRL1_mNG_ pMBS1239), mNeonGreen with mStop (mNG_Cyt_, pMBS1213), and mNeonGreen with PSAD transit peptide and mStop (mNG_cp_, pMBS1216), level 0 parts were directly assembled into level 2 destination vectors pMBS807 or pMBS808 already containing the *aadA* and *aphVIII* resistance cassettes, respectively (Niemeyer and Schroda, 2022). The level 0 parts of TurboID-C and the level 2 device for the mCherry_TID_ control were constructed previously (Kreis *et al*., 2023a).

### Protein analyses

Exponentially growing cells were harvested by centrifugation (5 min at 4,000 *g* and 4°C) and resuspended in DTT-SDS-sucrose buffer (0.1 M Na_2_CO_3_, 0.1 M DTT, 5 % (w/v) SDS, 30% (w/v) sucrose), boiled for 2 min at 95°C, and centrifuged at 12,000 *g* and 25°C for 5 min. Samples were subjected to SDS-PAGE and semi-dry western blotting based on chlorophyll concentrations (Porra *et al*., 1989). Antisera used were against the HA epitope (Sigma-Aldrich H3663, 1:10,000), BirA (Kreis *et al*. (2023a), 1:5,000), HSP70B (Drzymalla *et al*. (1996), 1:10,000), DEG1C (Theis *et al*. (2019) 1:5,000), HSP22E/F (Rütgers *et al*. (2017), 1:5,000), CLPB3 (Kreis *et al*. (2023b), 1:5,000), Cf1β (Lemaire and Wollman (1989), 1:5,000), VIPP2 (Nordhues *et al*. (2012), 1:5,000), CGE1 (Schroda *et al*. (2001), 1:3,000), Cyt *f* (Agrisera AS06 119, 1:5,000) or mNeonGreen (Agrisera AS21 4525, 1:5,000). Anti-rabbit-HRP (Sigma-Aldrich) and anti-mouse-HRP (Santa Cruz Biotechnology sc-2031) were used as secondary antibodies (1:10,000). Proteins were separated on a 12% SDS-PAGE gel and transferred to a nitrocellulose membrane, stained with Ponceau S (5 minute in 0.1% w/v Ponceau S in 5% acetic acid/water) and blocked with “blocking buffer” (3% w/v milk powder and 0.1% Tween - 20 in PBS) or “biotin blocking buffer” (3% w/v BSA and 0.1% Tween-20 in PBS) at 22°C for 30 min. The blots were incubated with the antisera in “blocking buffer” over-night at 4°C, rinsed 3 times with PBS-T (0.1% Tween20 in PBS) for 5 min and then incubated with the secondary antibody for at least 1 h at 22°C. For biotinylated proteins, the blots were immersed with streptavidin-HRP (Abcam ab7403, 1:20,000,) in “biotin blocking buffer” over-night at 4°C, then rinsed 3 times with PBS-T for 5 min. Immunodetection was performed using enhanced chemiluminescence (ECL) and the FUSION-FX7 Advance™imaging system (PEQLAB) or ECL ChemoStar V90D+ (INTAS Science Imaging).

### Crude fractionation

Exponentially growing cells were collected by centrifugation. The pellet was resuspended in 1 ml lysis buffer (10 mM Tris-HCl pH 8.0, 1 mM EDTA), 1% of protease inhibitor (Roche) was added and cells were transferred to a 2 mL Eppendorf tube. A 200 µl sample was taken as whole cell protein sample and supplemented with 70 µl 4x SDS buffer (10% SDS, 300 mM Tris HCl pH 8, 40% w/v sucrose, 0.05% Bromophenol-blue) and 50 µl of 1 M DTT. 200 µl of the remaining cells were fractionated by 4 cycles of freezing in liquid nitrogen and thawing at 25°C. After centrifugation for 20 min at 21,000 *g* and 4°C, 200 µl of the supernatant were taken as the soluble protein sample and supplemented with 70 µl of 4x SDS and 50 µl of 1 M DTT. The remaining pellet was resuspended in 200 µl lysis buffer, vortexed, and supplemented with 4x SDS buffer and 1M DTT as before to obtain the membrane protein sample. Samples were heated for at least 2 min at 95°C and briefly centrifuged at 18,500 *g* at 25°C before subjecting to SDS-PAGE for immunodetection.

### *In vivo* biotinylation, streptavidin affinity purification, MS sample preparation and mass spectrometry

For TurboID-mediated proximity-labeling, *in vivo* biotin labeling and streptavidin affinity purification, as well as MS sample preparation were carried out as described previously (Kreis *et al*., 2023a). Mass spectrometry on an LC-MS/MS system (Eksigent nanoLC 425 coupled to a TripleTOF 6600, ABSciex) was performed as described previously (Hammel *et al*., 2018). In brief: For peptide separation, an HPLC flow rate of 4 μl/min was used and gradients (buffer A 2% acetonitrile, 0.1% formic acid; buffer B 90% acetonitrile, 0.1% formic acid) ramped within 48 min from 3% to 35% buffer B, then to 50% buffer B within 5 min and finishing with wash and equilibration steps. MS1 spectra (350 m/z to 1250 m/z) were recorded for 250 ms and 35 MS/MS scans (110 m/z to 1600 m/z) were triggered in high sensitivity mode with a dwell time of 60 ms resulting in a total cycle time of 2,400 ms. Analyzed precursors were excluded for 10 s, singly charged precursors or precursors with a response below 150 cps were excluded from MS/MS analysis.

### Evaluation of MS data

Sciex raw data were converted to mzML format using MSConvert (Adusumilli and Mallick, 2017). The peptide-spectra-matching, protein inference and quantitation of the MS runs was performed using the Fragpipe v22.0 pipeline, integrating MSFragger, IonQuant, Percolator and Philosopher tools (da Veiga Leprevost *et al*., 2020; Käll *et al*., 2007; Kong *et al*., 2017; Yang *et al*., 2023; Yu *et al*., 2020). *Chlamydomonas reinhardtii* genome release 6.1 (Merchant *et al*., 2007) including chloroplast and mitochondrial proteins as well as TurboID and mCherry sequences and common contaminants were used as search space. Oxidation of methionine, biotinyl-lysine and acetylation of the N-terminus were considered as peptide modifications. Maximal missed cleavages were set to 3 and peptide length to 6 amino acids. Thresholds for peptide spectrum matching and protein identification were set by a false discovery rate (FDR) of 0.01. The mass spectrometry proteomics data have been deposited to the DATAPLANT XXX repository with the dataset identifier doi.:XXXXXX.

### MS data analysis – identification of enriched proteins

MS data analysis was carried out as described previously (Kreis *et al*., 2023a). Filtering and statistical analysis were done with Perseus (version 1.6.15.0) (Tyanova *et al*., 2016). The ‘combined.protein.tsv’ output file from MSFragger/FragPipe (Kong *et al*., 2017) was imported into Perseus using the intensity values as main category and matched with the Metadata protein annotation v6.1 of *Chlamydomonas reinhardtii.* The data matrix was filtered to remove contaminants and reverse measurements. Intensity values were log2 transformed and proteins were kept that were identified/quantified in three out of four replicates of at least one strain. Normalization was achieved by subtracting the median using columns as matrix access. Missing values were imputed for statistical analysis using the ‘Replace missing values from normal distribution’ function with the following settings: width = 0.3, down shift = 1.8, and mode = total matrix. To identify proteins significantly enriched in T_ID_-expressing samples versus the mCherry_TID_ control, unpaired two-tailed Student’s t-tests were performed comparing CDJ5-WT_TID_, CDJ5-AAA_TID_, CDJ5-SSS_TID_, LCP1_TID_, QCP1_TID_, and PGRL1_TID_ with the mCherry_TID_#4 samples. The integrated modified permutation-based FDR was used for multiple sample correction with an FDR of 0.05, an S0 of 1, and 250 randomizations to determine the cutoff. Significantly enriched proteins were kept. Finally, to determine proteins that were enriched compared to mCherry_TID_, the cutoff parameters were set to a q-value ≤ 0.05 and a minimal 4- fold enrichment. Volcano plots were made in Excel using t-test results exported from Perseus.

### Microscopy

Light and fluorescence microscopy images were taken with an Olympus BX53 microscope. Fluorescent signals from fluorescein-isothiocyanate (FITC; excitation/emission 494 nm/518 nm), mNeonGreen and mCherry (excitation/emission 580 nm/610 nm), and chlorophyll autofluorescence were recorded with the following parameters: lamp intensity 100, gain 1.5 for 150-600 ms. Signals were processed using ImageJ (Fiji plug-in).

### Computational analyses

Predictions of subcellular localization was performed with PredAlgo (Tardif *et al*., 2012), TargetP (Almagro Armenteros *et al*., 2019), Plant-mSubP (Sahu *et al*., 2019), BacCelLo (Pierleoni *et al*., 2006), PB-Chlamy (Wang *et al*., 2023), and SUBAcon (Hooper *et al*., 2014). Structures of LCP1 and QCP1 were predicted by AlphaFold2 (Jumper *et al*., 2021) via ColabFold (Mirdita et al. 2022). Pairwise structural comparisons were done with the Analyze tool in RCSB (https://www.rcsb.org/) (Berman *et al*., 2000) and displayed with Mol* (Sehnal *et al*., 2021). Sequence alignments were done with CLUSTALW (https://www.genome.jp/tools-bin/clustalw) and displayed with GeneDoc.

## Results

The aim of this study was to employ TurboID-based proximity labeling to identify proteins in the proximity of CDJ5. Since we had established TurboID as a genetic part for the Modular Cloning (MoClo) strategy (Kreis *et al*., 2023a), we first had to generate a MoClo-compatible genetic part encoding wild-type (WT) CDJ5. For this, we synthesized the *CDJ5* cDNA interrupted by the three introns of the *Chlamydomonas RBCS2* gene to enhance transgene expression (CDJ5-WT) (Baier *et al*., 2018; Schroda, 2019). We next changed the codons for histidine, proline and aspartate of the HPD motif in the J domain to alanine codons (CDJ5- AAA) and three of the four codons for cysteines that mediate the attachment of the 4Fe-4S cluster to the polypeptide chain to serine codons (CDJ5-SSS). The rationale for mutating the HPD motif was that we expected substrate proteins bound by CDJ5 to remain associated with CDJ5 for a longer time when their transfer to its HSP70 partner is perturbed by the impaired stimulation of HSP70’s ATPase activity. The rationale for the mutant unable to bind a 4Fe-4S cluster was that we expected this variant not to interact with substrate proteins requiring a functional 4Fe-4S cluster. Hence, proteins identified in the proxiome of CDJ5-WT and CDJ5- AAA but not in that of CDJ5-SSS likely depend on the 4Fe-4S cluster for interacting with CDJ5.

### CDJ5 localizes to the chloroplast and there mainly to the soluble fraction but also to the membrane-enriched fraction

To define a proper control for proximity labeling with CDJ5, we first had to get information on its intracellular localization. Since all our attempts to produce CDJ5 recombinantly in *E. coli* for raising a polyclonal antibody had failed, we needed to express tagged CDJ5 in *Chlamydomonas*. For this, we assembled two transcriptional units containing the genetic part encoding CDJ5-WT, parts containing the strong *HSP70A-RBCS2* (*AR*) promoter and *RPL23* terminator (Lopez-Paz *et al*., 2017; Schroda *et al*., 2000) as well as parts encoding a 3xHA tag or mNeonGreen (mNG). The two transcriptional units were then combined with one that drives expression of the *aadA* cassette (S^R^) (Fig. 1A). We readily obtained spectinomycin resistant transformants producing CDJ5-WT_3xHA_ and CDJ5-WT_mNG_ that migrated on SDS-PAGE with the expected molecular masses of 41.6 and 64.5 kDa (after removal of the predicted chloroplast transit peptide (Dorn *et al*., 2010)) (Fig. 1B, D). Fractionation experiments performed with the transformant producing CDJ5-WT_3xHA_ revealed the protein to localize mainly in the soluble fraction; however, a substantial part of CDJ5_3xHA_ was also detected in the membrane-enriched pellet fraction (Fig. 1B, C). CDJ5-WT fused to mNG allowed localizing the protein in the chloroplast in comparison to free mNG targeted to the chloroplast stroma via the transit peptide of PSAD and to the cytosol (Figs. 1A, 1E; Supplementary Figs. S1-S3). CDJ5-WT_mNG_ fluorescence co-localized mostly with chlorophyll fluorescence while stromal mNG fluorescence mainly originated from cavities surrounded by Chl fluorescence. This corroborates the previous localization of CDJ5 as “enriched in thylakoid” (Wang *et al*., 2023). Moreover, CDJ5-WT_mNG_ fluorescence was enriched in the pyrenoid, where in most images it covered the entire pyrenoid region. In some images more delicate structures were discernible that could correspond to pyrenoid tubules (Mackinder *et al*., 2017; Wang *et al*., 2023). In contrast, stromal mNG gave rise to an intense fluorescence spot in the middle of the pyrenoid which was surrounded by a ring with low fluorescence.

**Fig. 1.**
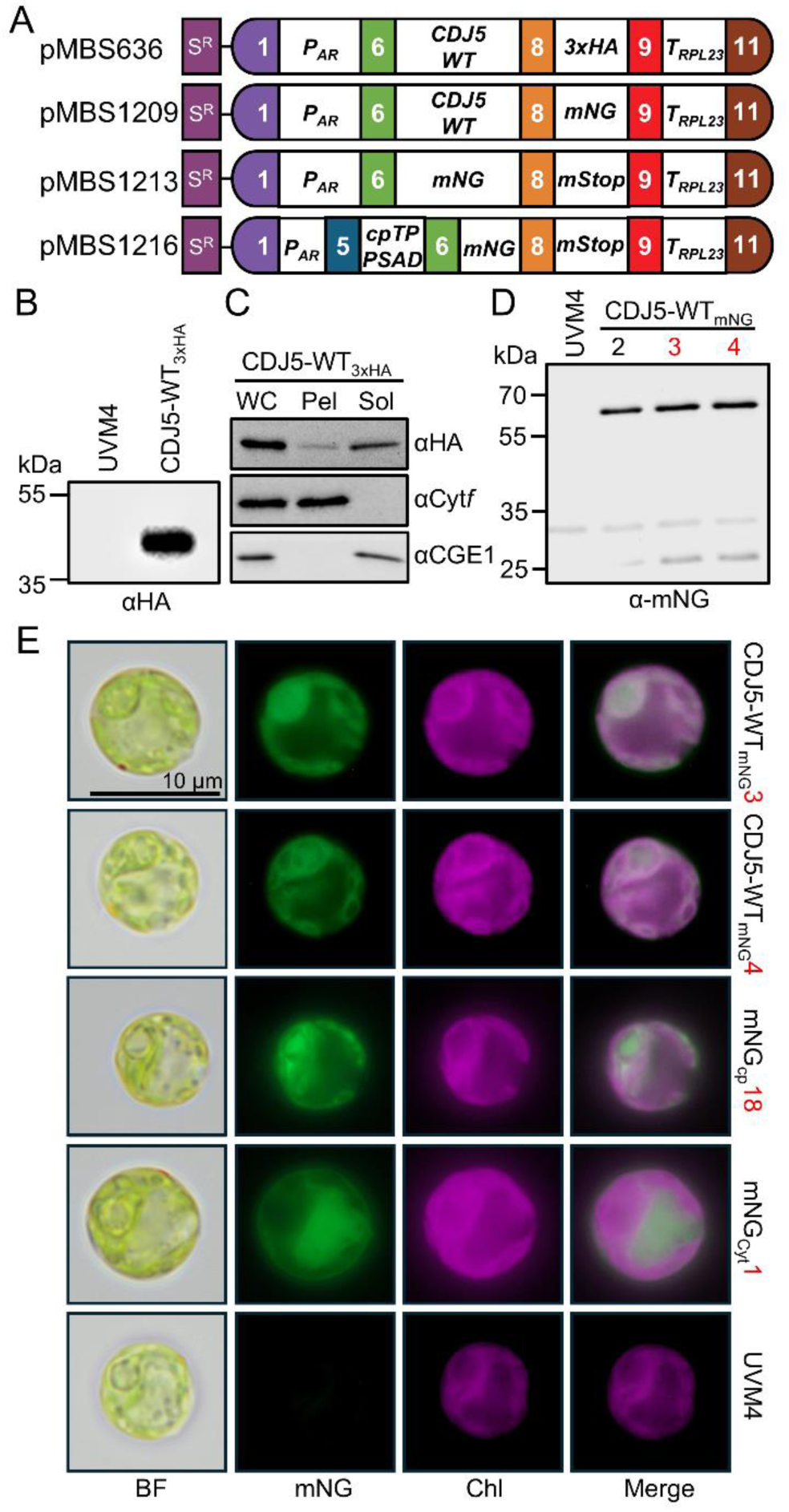
Overexpression of CDJ5-WT and localization. (A) Level 2 constructs conferring resistance to spectinomycin (S^R^) and driving the production of CDJ5-WT with C-terminal 3xHA or mNeonGreen (mNG) tags as well as of free mNG targeted to the cytosol and the stroma via the PSAD chloroplast transit peptide (cpTP_PSAD_). Gene expression is controlled by the *HSP70A-RBCS2* promoter (*P_AR_*) and the *RPL23* terminator (*T_RPL23_*). The mStop part contains three stop codons. (B) Analysis of CDJ5-WT_3xHA_ protein accumulation. Total cell proteins from a transformant generated with pMBS636 and from the UVM4 recipient strain were extracted, separated by SDS-PAGE (≥2 µg chlorophyll) and analyzed by immunoblotting using an antibody against the HA-epitope. (C) Crude fractionation of cells producing CDJ5-WT_3xHA_. Whole cell proteins (WC), pellet (Pel) and soluble (Sol) fractions generated by three freeze/thaw cycles and centrifugation were separated by SDS-PAGE and analyzed by immunoblotting using antibodies against soluble stromal CGE1, membrane-integral Cyt*f* and the HA-epitope. (D) Analysis of CDJ5-WT_mNG_ accumulation. Total cell proteins from three transformants generated with pMBS1209 and from the UVM4 recipient strain were extracted, separated by SDS-PAGE (2 µg chlorophyll) and analyzed by immunoblotting using an antibody against mNG. Transformants labeled with red numbers were used for subsequent localization studies. (E) Localization of CDJ5-WT_mNG_ and of free mNG targeted to the stroma (mNG_cp_) and the cytosol (mNG_Cyt_). UVM4 cells served as negative control. Shown are bright field (BF) images, mNG fluorescence (green) and chlorophyll autofluorescence (Chl, magenta). Merge – overlap of both fluorescence signals. The scale bar applies to all images.

### Proximity labeling reveals HSP70B to be close to all three CDJ5 variants

We next assembled three transcriptional units utilizing the genetic parts encoding CDJ5-WT, CDJ5-AAA, and CDJ5-SSS, parts containing the *HSP70A-RBCS2* promoter and the *RPL23* terminator as well as the part encoding TurboID (T_ID_) (Kreis *et al*., 2023a). The resulting transcriptional units were combined with the one driving expression of the *aadA* cassette (S^R^) (Fig. 2A). We readily obtained spectinomycin resistant transformants that produce the three CDJ5_TID_ variants migrating on SDS gels with the expected molecular mass of 73.6 kDa (after removal of the predicted chloroplast transit peptide) (Fig. 2B). We chose to use stromally targeted mCherry_TID_ as control for proximity labeling, which we had employed previously as control for VIPP1/2_TID_ (Kreis *et al*., 2023a). We used two control strains (#13 and #4) that accumulated mCherry_TID_ to lower (#13) and higher levels (#4).

**Fig. 2.**
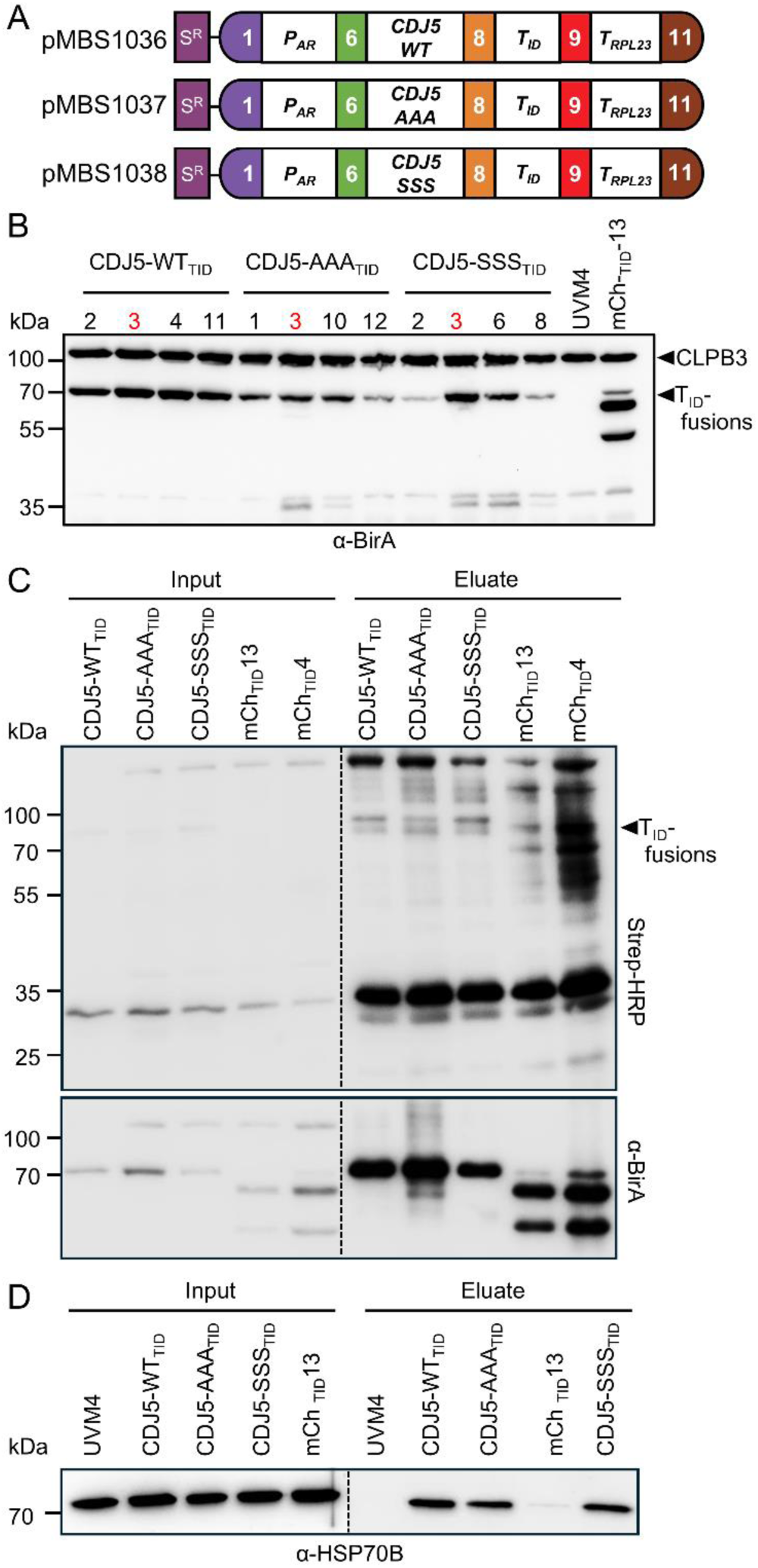
TurboID-based proximity labeling using CDJ5 variants as baits. (A) Level 2 constructs conferring resistance to spectinomycin (S^R^) and driving the production of CDJ5-WT, CDJ5-AAA or CDJ5-SSS with C-terminal TurboID fusion (T_ID_). Gene expression is controlled by the *HSP70A-RBCS2* promoter (*P_AR_*) and the *RPL23* terminator (*T_RPL23_*). (B) Analysis of CDJ5-WT_TID_, CDJ5-AAA_TID_ and CDJ5-SSS_TID_ protein accumulation. Total cell proteins were extracted from four transformants for each construct shown in (A), separated by SDS-PAGE (2 µg chlorophyll) and analyzed by immunoblotting using an antibody against BirA. UVM4 served as negative control, mCh_TID_-13 with TurboID fused to mCherry as positive control. Transformants numbered in red were chosen for proximity labeling experiments. The positions of the TurboID fusion proteins and CLPB3, cross-reacting with the antibody, are indicated. (C) Enhancement of TurboID *in-vivo* biotinylation activity by externally added biotin. Cultures of transformants producing the CDJ5_TID_ variants, UVM4 (sample not loaded here) and two transformants producing mCh_TID_ were grown at 25°C to mid-log phase and supplemented with 1 mM biotin for 4 h. Cells were harvested and lysed (input) whereafter biotinylated proteins were enriched by affinity purification with streptavidin beads (eluate). Proteins were separated by SDS-PAGE and analyzed by immunoblotting with streptavidin-HRP and with an antibody against BirA. One of four biological replicates is shown. (D) Verification of HSP70B in the vicinity of CDJ5. Samples from (C) were separated by SDS-PAGE and analysed by immunoblotting with an antibody against HSP70B.

The three CDJ5_TID_ variants, the two mCherry_TID_ controls and the UVM4 recipient strain were grown to mid-log phase and supplemented with 1 mM biotin for 4 h, corresponding to the optimal labeling conditions for chloroplast TurboID in *Chlamydomonas* (Kreis *et al*., 2023a; Lau *et al*., 2023). The detection of biotinylated proteins in whole-cell protein extracts with streptavidin-horseradish peroxidase (HRP) revealed hardly any self-biotinylation of the three CDJ5_TID_ variants and the mCherry_TID_ controls. However, all five bait proteins were strongly enriched after purification with streptavidin beads, as detected with streptavidin-HRP and the antibody against TurboID (αBirA) (Fig. 2C). Immunodetection of HSP70B revealed that it was not present in eluates from streptavidin beads incubated with protein extracts of the UVM4 recipient strain, weakly present in eluates from the mCherry_TID_ control line #13 and clearly present in eluates from transformants producing the three CDJ5_TID_ variants (Fig. 2D). The weak presence of HSP70B in eluates from the mCherry_TID_ control is expected since HSP70B is an abundant protein (∼0.19% of total cell proteins (Liu *et al*., 2007)) and should become non-specifically biotinylated to some extent by stromal mCherry_TID_. The clear presence of HSP70B in eluates from all three CDJ5_TID_-producing transformants suggested that the recombinant CDJ5_TID_ variants are generally functional and that the mutated HPD motif in CDJ5-AAA and the absence of the 4Fe-4S cluster in CDJ5-SSS do not interfere with the ability of these variants to interact with HSP70B.

### The proxiomes of CDJ5-WT and CDJ5-AAA largely overlap but that of CDJ5-SSS is very different

We analysed the streptavidin eluates from four independent experiments performed as shown in Fig. 2C by mass spectrometry using a TripleTOF 6600 instrument. Data analysis using MSFragger followed by Perseus yielded 1366 protein groups. Here proteins were kept that were identified/quantified in three out of four replicates of at least one strain (Supplementary Dataset S1). The volcano plots shown in Fig. 3A-C show the contrasts between the respective CDJ5 variant and the mCherry_TID_#4 control. 85 proteins were significantly enriched (≥ 2 peptides, > 4-fold, q ≤ 0.05) with CDJ5-AAA_TID_, whereas 30 and 37 proteins were enriched with CDJ5-WT_TID_ and CDJ5-SSS_TID_, respectively (excluding the CDJ5_TID_ baits). 88% of the proteins in the CDJ5-AAA proxiome, all proteins in the CDJ5-WT proxiome, and 95% in the CDJ5-SSS proxiome were predicted to be chloroplast-localized by at least one of six prediction programs used (Supplementary Dataset 1). The larger number of proteins enriched with CDJ5-AAA_TID_ could be due to a longer dwelling time of substrates on this CDJ5 variant since their transfer to HSP70B should be impaired by the mutated HPD motif. However, it could also be due to the higher expression level of CDJ5-AAA_TID_ compared to CDJ5-WT_TID_ and CDJ5-SSS_TID_ (Fig. 2C).

**Fig. 3.**
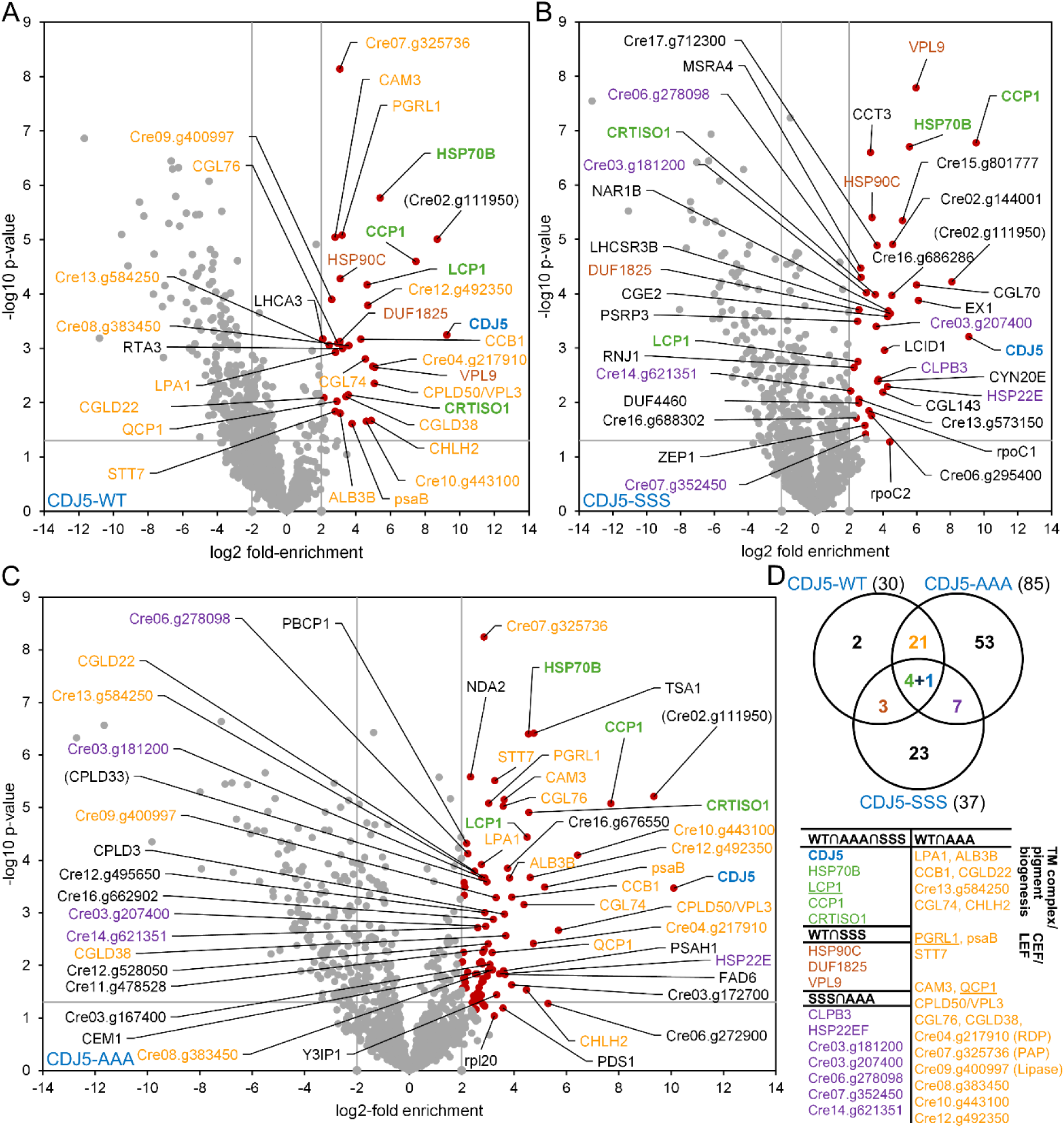
Proximity labeling results for the three CDJ5 variants. (A-C) Volcano plots of the proxiomes of CDJ5-WT, CDJ5-SSS, and CDJ5-AAA. Streptavidin bead eluates of the samples shown in Fig. 2C were analyzed by mass spectrometry. Shown is the log2-fold enrichment between proteins in the CDJ5_TID_ and mCh_TID_#4 proxiomes. Proteins significantly enriched in the CDJ5_TID_ samples (q ≤ 0.05, enrichment ≥ 4) are shown as red data points. In the CDJ5-AAA dataset, proteins with enrichment between 4 and 8-fold and a q value between 0.01 and 0.05 are not labeled to reduce complexity. The CDJ5 bait is shown in blue, proteins enriched with all three CDJ5 variants in green, proteins enriched with CDJ5-WT and CDJ-AAA in orange, proteins enriched with CDJ5-WT and CDJ5-SSS in brown, and proteins enriched with CDJ5-SSS and CDJ5-AAA in purple. Cre02.g111950 and CPLD33 were identified with a single peptide only and are therefore in parentheses. (D) Summary of results. The Venn diagram shows the number of proteins enriched with each CDJ5 variant and their overlap. The table summarizes the IDs of the enriched proteins. Proteins found in all three proxiomes are not listed again in the overlaps of two proxiomes.

The CDJ5_TID_ baits were enriched between 544- and 1100-fold. We found four proteins significantly enriched with all three baits, which are HSP70B (23-48-fold); transporter CCP1 (177- to 735-fold) which with its paralog CCP2 is required for growth in a low-CO_2_ environment (Pollock *et al*., 2004); CRTISO1 (12- to 24-fold), which is involved in carotenoid biosynthesis (Lohr, 2023); and Lon domain-containing protein 1 (LCP1, see below) (5.7- to 25-fold) (Fig. 3, green letters; Supplementary Dataset S1). HSP90C, a chaperone found in complex with HSP70B (Willmund *et al*., 2008), was also significantly enriched up to 10.2-fold with all three CDJ5 variants but in case of CDJ5-AAA remained just below the 4-fold enrichment cut-off (3.97-fold enriched). Strikingly, 25 of the 30 proteins in the CDJ5-WT proxiome were shared by the CDJ5-AAA proxiome, whereas CDJ5-WT and CDJ5-SSS shared only 7 proteins in their proxiomes. Similarly, CDJ5-AAA and CDJ5-SSS shared only 11 proteins in their proxiomes. These results suggest that the local environment of CDJ5 changes substantially when it is unable to bind a 4Fe-4S cluster. The significant enrichment of the disaggregase CLPB3 and the small heat shock proteins HSP22E/F only in the proxiomes of CDJ5-SSS (13- and 19-fold enrichment, respectively) and CDJ5-AAA (8- and 11-fold enrichment, respectively) suggested either that the introduced point mutations could have rendered the proteins more prone to aggregation, or could have triggered an unfolded protein response (cpUPR) (Gabelmann and Schroda, 2025). To test the latter scenario, we analysed the accumulation of cpUPR marker proteins CLPB3, DEG1C, VIPP2, and HSP22E/F in the lines producing the CDJ5 variants harbouring C-terminal 3xHA or TurboID fusions compared to the UVM4 strain treated with H_2_O_2_ for 4 h to trigger a cpUPR (Kreis *et al*., 2023a; Theis *et al*., 2020). As shown in Supplementary Fig. S4, none of the cpUPR markers was substantially upregulated in the CDJ5 variants when compared to the H_2_O_2_-treated control, indicating no negative effects of expressing the CDJ5-SSS and CDJ5-AAA variants on chloroplast proteostasis.

### Proteins involved in thylakoid membrane protein complex assembly, pigment biosynthesis, and regulation of photosynthetic electron flow are specifically enriched in the proxiomes of CDJ5 variants capable of binding a 4Fe-4S cluster

Of the 21 proteins present in the CDJ5-WT and CDJ5-AAA proxiomes but not in the CDJ5- SSS proxiome, five have functions in the biogenesis of thylakoid membrane protein complexes, i.e., LPA1 (∼7-fold enriched) and ALB3B (8.5- to 14-fold enriched) for the photosystems (Göhre *et al*., 2006; Peng *et al*., 2006); CCB1 (15- to 19.4-fold enriched) for the Cyt b_6_/f complex (Lezhneva *et al*., 2008); and CGLD22/CGL160 (4.4- to 5.6-fold enriched) and potentially Cre13.g584250 (5.5- to 7.4-fold enriched) for the ATP synthase (Reiter *et al*., 2022). Further proteins involved in thylakoid membrane protein complex biogenesis were enriched only in the CDJ5-AAA proxiome, i.e., Y3IP1 (10.1-fold enriched) and ycf3/pafI (7-fold enriched) for PSI biogenesis (Nellaepalli *et al*., 2021); and RBD1 (5.3-fold enriched) and HHL1 (6.6-fold enriched) for PSII biogenesis/repair (Garcia-Cerdan *et al*., 2019; Jin *et al*., 2014). In the context of thylakoid membrane protein complex biogenesis is the enrichment of proteins involved in chlorophyll biogenesis in the CDJ5-WT and CDJ5-AAA proxiomes. These are the subunit H2 of the Mg-chelatase 2 (CHLH2, 22.2-29.6-fold enriched (Chekounova *et al*., 2001) and protein CGL74 (20.8-23.4-fold enriched) (Herbst *et al*., 2024).

Three proteins enriched with CDJ5-WT and CDJ5-AAA play roles in driving/controlling photosynthetic electron flow: PGRL1 (8.2- to 9.2-fold enriched) and psaB (13.6- to 36-fold enriched) are part of a complex driving cyclic electron flow (CEF) (DalCorso *et al*., 2008; Iwai *et al*., 2010) and STT7 (7- to 9.6-fold enriched) regulates state transitions (Depege *et al*., 2003). Several more proteins identified only in the CDJ5-AAA or CDJ5-WT proxiomes also play roles in CEF, i.e., psaA (6.4-fold enriched), PSAH1 (12-fold enriched), LHCA3 (4.2-fold enriched), and CP26/LHCB5 (4.3-fold enriched) as components of the PGRL1-containing CEF complex (Iwai *et al*., 2010) and NDA2 (5-fold enriched) involved in an alternative CEF pathway (Saroussi *et al*., 2016). Furthermore, phosphatase PBCP1 (4.6-fold enriched) antagonizing state transition kinase (Cariti *et al*., 2020) was found in the CDJ5-AAA proxiome.

### LCP1, QCP1, and PGRL1 localize to distinct structures in the chloroplast

To exemplarily validate the proxiomes particularly of the CDJ5 variants capable of binding a 4Fe-4S cluster, we chose LCP1 as a protein enriched in the proxiomes of all three CDJ5 variants as well as PGRL1 and QCP1 as proteins found only in the CDJ5-WT and CDJ5-AAA proxiomes (Fig. 3). LCP1 consists only of the N domain of LonA AAA+ proteases while it lacks the central ATPase domain and the C-terminal proteolytic domain (Wlodawer *et al*., 2022). LCP1 is predicted to be chloroplast targeted, lacks transmembrane domains, is conserved in the green lineage and its AlphaFold2 structure is very similar to the crystal structure determined for the N domain of the Lon protease from *Mycobacterium avium* (RMSD = 3.07 A; TM score 0.78; Supplementary Fig. S5) (Chen *et al*., 2019). In Arabidopsis, LCP1 was shown to inhibit Trx-y2 resulting in an enhanced accumulation of H_2_O_2_ (Shin *et al*., 2020). PGRL1 contains two transmembrane helices and as mentioned is involved in CEF. PSBQ domain-containing protein 1 (QCP1) is predicted to be chloroplast targeted, lacks transmembrane domains, and is conserved in the green lineage up to mosses (Supplementary Fig. 6A). QCP1 contains a domain that is structurally very similar to the PSBQ protein of the oxygen evolving complex of PSII, as judged from a pairwise structural alignment of PSBQ from spinach and the AlphaFold2 structure of QCP1 (Supplementary Fig. S6B; RMSD = 2.07 A; TM score 0.81). However, the function of QCP1 is unknown.

We first wanted to localize the three selected proteins. For this, we synthesized codon-optimized sequences as MoClo level 0 parts encoding LCP1 and QCP1 interspersed once and twice with *RBCS2* intron 1, respectively, and once with *RBCS2* intron 2. The two gene parts and the one for *PGRL1* reported previously (Kreis *et al*., 2023a) were then assembled into a *Chlamydomonas* expression vector already containing the *aphVIII* paromomycin resistance cassette (P^R^) (Niemeyer and Schroda, 2022) with parts containing the *HSP70A-RBCS2* promoter and the *RPL23* terminator as well as the part encoding mNG. Since the *PGRL1* part lacked the sequence encoding its chloroplast transit peptide, we added a part encoding the PSAD transit peptide for *PGRL1* (Fig. 4A). We readily obtained paromomycin resistant transformants that produce the three protein variants migrating on SDS gels with the expected molecular masses of 54.7 kDa (LCP1), 56.5 kDa (QCP1), and 58.6 kDa (PGRL1) after removal of the predicted chloroplast transit peptide (Fig. 4B). Fluorescence microscopy revealed that PGRL1_mNG_ fluorescence largely co-localized with chlorophyll fluorescence. Moreover, PGRL1_mNG_ fluorescence gave rise to a small spot in the middle of the pyrenoid with delicate radial extensions presumably deriving from thylakoid membrane tubules (Fig. 4C; Supplementary Fig. S7). Fluorescence from LCP1_mNG_ also co-localized largely with chlorophyll fluorescence and was present in delicate structures in the middle of the pyrenoid (Fig. 4C; Supplementary Fig. S8). Finally, fluorescence from QCP1_mNG_ again co-localized largely with chlorophyll fluorescence while bright fluorescence was again present in the pyrenoid region (Fig. 4C; Supplementary Fig. S9). In contrast to CDJ5_mNG_, PGRL1_mNG_, and LCP1_mNG_, we always observed mNG fluorescence in QCP1_mNG_ transformants also in the cytosol. We detected an additional protein band of ∼25 kDa in QCP1_mNG_ and CDJ5_mNG_ transformants that cross-reacted with the mNG antibody and co-migrated in SDS gels with free mNG (Fig. 1D; Supplementary Figs. 2B, 3B). Since we observed mNG fluorescence only in the cytosol of QCP1_mNG_ transformants but not in CDJ5_mNG_ transformants, we assume that mNG was cleaved off from QCP1_mNG_ prior to entering the chloroplast, while it was cleaved off from CDJ5_mNG_ after entering the chloroplast.

**Fig. 4.**
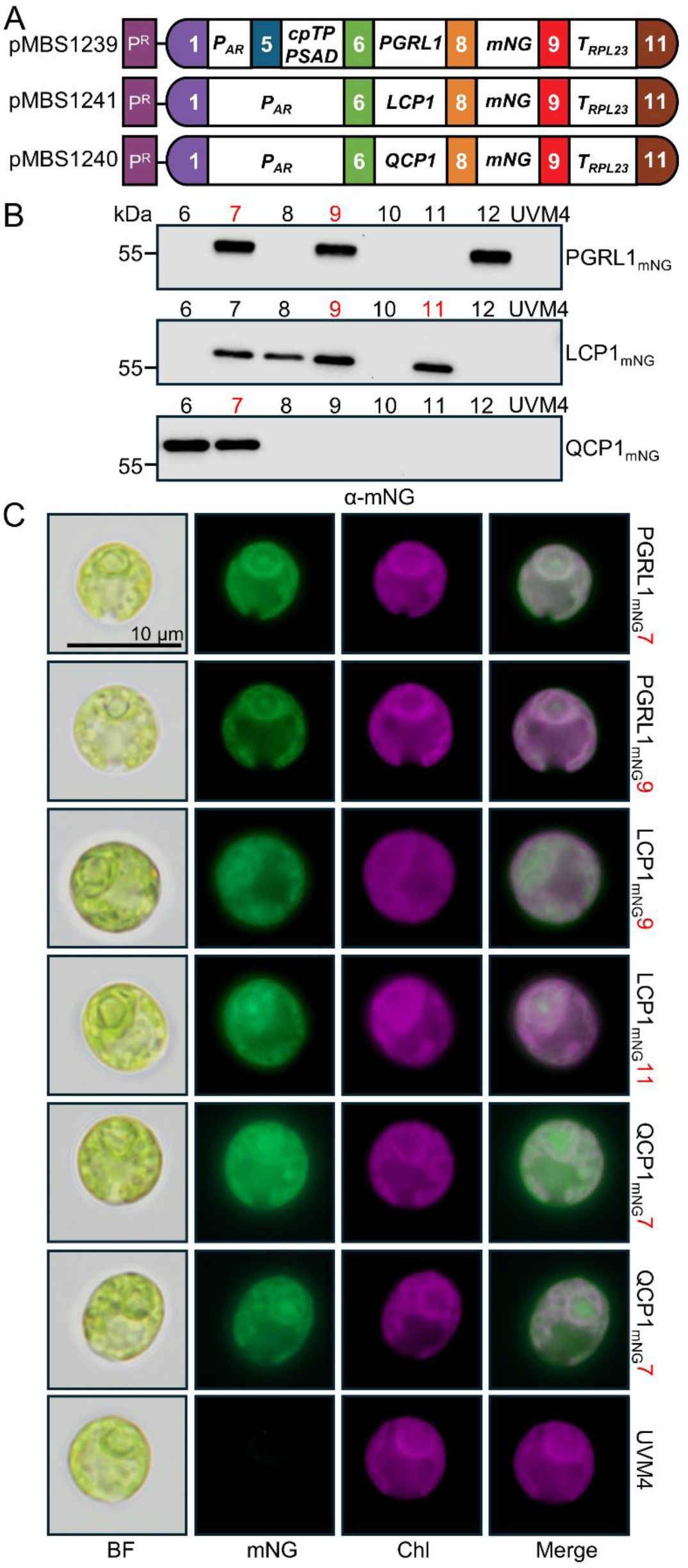
Localization of CDJ5-proximal proteins. (A) Level 2 constructs conferring resistance to paromomycin (P^R^) and driving the production of PGRL1, LCP1, and QCP1 with C-terminal mNeonGreen (mNG) tag. Gene expression is controlled by the *HSP70A-RBCS2* promoter (*P_AR_*) and the *RPL23* terminator (*T_RPL23_*). PGRL1 contains the chloroplast transit peptide (cpTP) of PSAD. ( B) Analysis of mNG fusion protein accumulation. Total cell proteins from transformants generated with the constructs in (A) and from the UVM4 recipient strain were extracted, separated by SDS-PAGE (2 µg chlorophyll) and analyzed by immunoblotting using an antibody against the mNG. Transformants labeled in red were chosen for microscopy. (C) Localization of mNG fusions to PGRL1, LCP1, and QCP1 and UVM4 as control. Shown are bright field (BF) images, mNG fluorescence (green) and chlorophyll autofluorescence (Chl, magenta). Merge – overlap of both signals. The scale bar applies to all images.

### CDJ5 is present in the proxiomes of LCP1 and QCP1 but not in that of PGRL1

To validate the proximity of CDJ5 to LCP1, QCP1, and PGRL1 we assembled level 2 constructs for the expression of LCP1 and QCP1 with C-terminal TurboID fusions under control of the *HSP70A-RBCS2* promoter and the *RPL23* terminator. The parts were directly assembled into a vector already containing the *aphVIII* paromomycin resistance cassette (P^R^) (Niemeyer and Schroda, 2022) (Fig. 5A) and transformed into the strain expressing CDJ5- 3xHA. Construct and transgenic line expressing PGRL1 with PSAD chloroplast transit peptide under control of the *PSAD* promoter and *RPL23* terminator are described in Kreis et al. (2023). Paromomycin resistant transformants producing the LCP1- and QCP1-TurboID fusions were readily obtained (Fig. 5B). Both proteins migrated in SDS-PAGE with the expected molecular masses of 63.9 kDa (LCP1_TID_) and 65.6 kDa (QCP1_TID_), however, in the QCP1_TID_ strain we detected an additional protein band migrating at ∼35 kDa that cross-reacted with the BirA antibody and most likely represents TurboID cleaved off from the QCP1_TID_ fusion protein at the linker sequence. Addition of biotin to the cultures for 4 h and subsequent streptavidin affinity purification resulted in the enrichment of all three TurboID fusion proteins, as detected with streptavidin-HRP and the antibody against TurboID (αBirA) (Fig. 5C). CDJ5-WT_3xHA_ was detected in the input samples from lines expressing PGRL1_TID_, LCP1_TID_ and QCP1_TID_ and in the streptavidin eluates only in the lines expressing QCP1_TID_ and LCP1_TID_ but not in that expressing PGRL1_TID_.

**Fig. 5.**
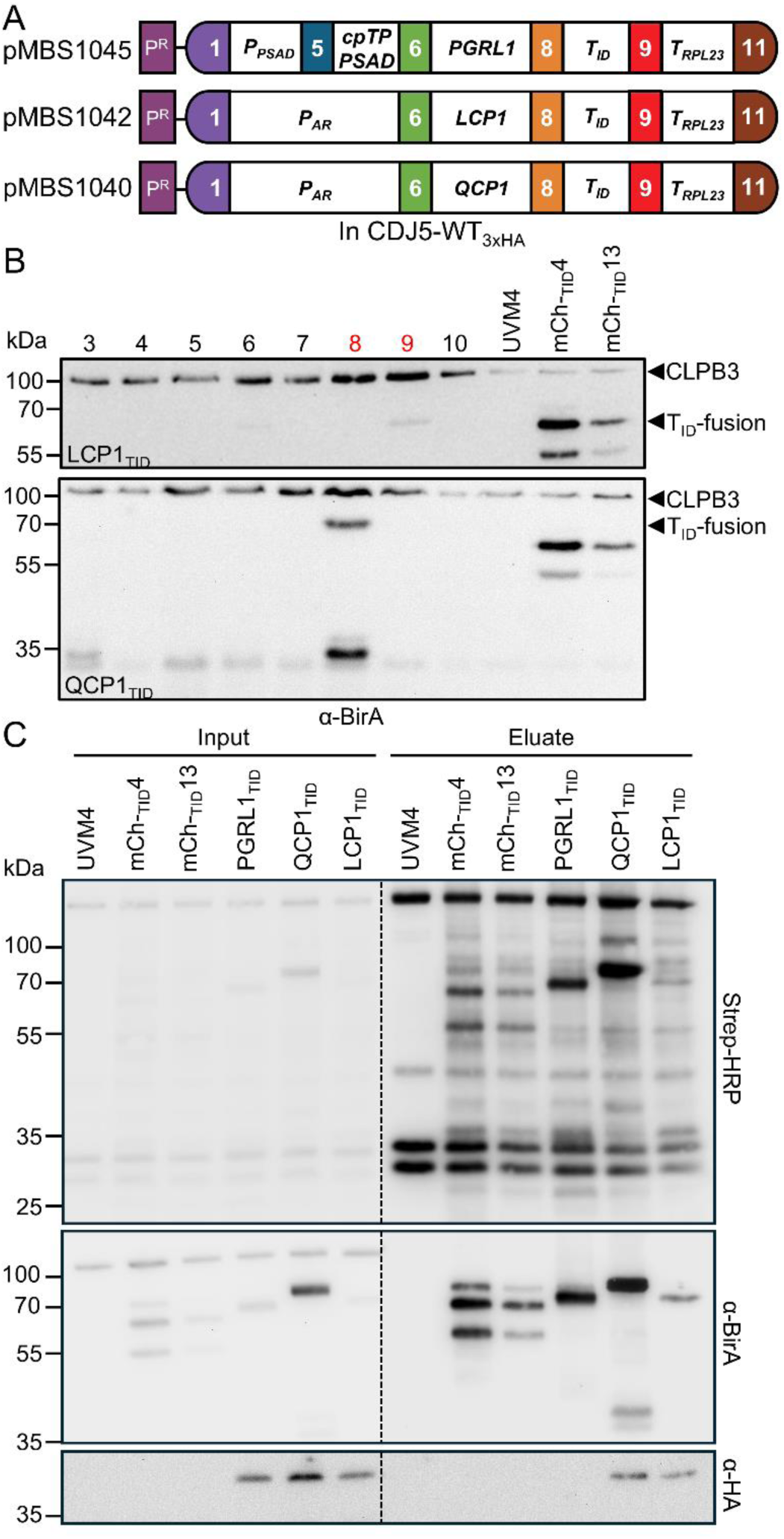
Reciprocal TurboID-based proximity-labeling using PGRL1, LCP1 and QCP1 as baits. (A) Level 2 constructs conferring resistance to paromomycin (P^R^) and driving the production of PGRL1, LCP1 and QCP1 with C-terminal TurboID fusion (T_ID_). Gene expression is controlled by the *PSAD* promoter (*P_PSAD_*) or the *HSP70A-RBCS2* promoter (*P_AR_*), and the *RPL23* terminator (*T_RPL23_*). The fusion protein of PGRL1 is targeted to the chloroplast via the *PSAD* chloroplast transit peptide (*cpTP PSAD*). Plasmids were transformed into the strain producing CDJ5-WT_3xHA_ (Figure 1B). (B) Analysis of LCP1_TID_ and QCP1_TID_ protein accumulation. Total cell proteins were extracted from eight transformants for constructs pMBS1042 and pMBS1040 shown in (A), separated by SDS-PAGE (2 µg chlorophyll) and analyzed by immunoblotting using an antibody against BirA. UVM4 served as negative control, mCh_TID_-4 and 13 with TurboID fused to mCherry as positive controls. Transformants numbered in red were chosen for proximity labeling experiments. The positions of the TurboID fusion proteins and CLPB3, cross-reacting with the BirA antibody, are indicated. (C) TurboID-mediated *in-vivo* biotinylation after addition of biotin. Cultures of transformants producing PGRL1_TID_, LCP1_TID_ and QCP1_TID_, UVM4, and mCh_TID_-4 and 13 were grown at 25°C to mid-log phase and supplemented with 1 mM of biotin for 4 h. Cells were harvested and lysed (input) whereafter biotinylated proteins were enriched by affinity purification with streptavidin beads (eluate). Proteins were separated by SDS-PAGE and analyzed by immunoblotting with an antibody against BirA and the HA epitope (to detect CDJ5-WT_3xHA_), and with streptavidin-HRP. One of four biological replicates is shown.

### The proxiomes of LCP1, QCP1 and PGRL1 share only few proteins

We analysed the streptavidin eluates from four independent experiments performed as shown in Fig. 5C with the three strains and the mCherry_TID_#4 control by mass spectrometry and using MSFragger followed by Perseus identified 1117 protein groups (Supplementary Dataset S2). The volcano plots shown in Fig. 6A-C show the contrasts between LCP1_TID_, QCP1_TID_, and PGRL1_TID_ and the mCherry_TID_#4 control. 34 proteins were significantly enriched with LCP1_TID_ (excluding LCP1) (Fig. 6A), 88% of which were predicted to be localized to the chloroplast by at least one of six prediction programs (Supplementary Dataset S2). LCP1_TID_ was enriched only 97-fold, presumably because it was only weakly expressed (Fig. 5B, C). CDJ5 was enriched 12.4-fold. Of the proteins found in the proxiomes of the CDJ5 variants, only five proteins in addition to CDJ5 were also found in the LCP1 proxiome: these are CRTISO1 found with all CDJ5 variants (13.5-fold enriched); AKC4/VPL7 and Cre02.g108250 found with CDJ5- WT and CDJ5-AAA (6.4- and 7-fold enriched, respectively); and EX1 and CYN20E found with CDJ5-SSS (315- and 7.9-fold enriched, respectively). In addition to the extreme enrichment of EX1, the ortholog of the Arabidopsis EXECUTER proteins (Wagner *et al*., 2004), it is remarkable that two AAA+ ATPases were found in the LCP1 proxiome, which are HSLU2 (75.1-fold enriched) and CLPC1 (4.7-fold enriched) as well as clpP as a subunit of the connected proteolytic chamber (4.8-fold enriched) (Schroda and deVitry, 2023). Other proteins in the LCP1 proxiome with common functions are translation factors TDA1 (*atpA* translation, 42.6-fold enriched) (Eberhard *et al*., 2011) and TAB2 (*psaB* translation, 6.9-fold enriched) (Dauvillee *et al*., 2003) as well as ACS2, PDH2, and KAS1 involved in fatty acid biosynthesis (enriched 5.5-, 4.6-, and 9.9-fold, respectively) (Li-Beisson *et al*., 2023).

**Fig. 6.**
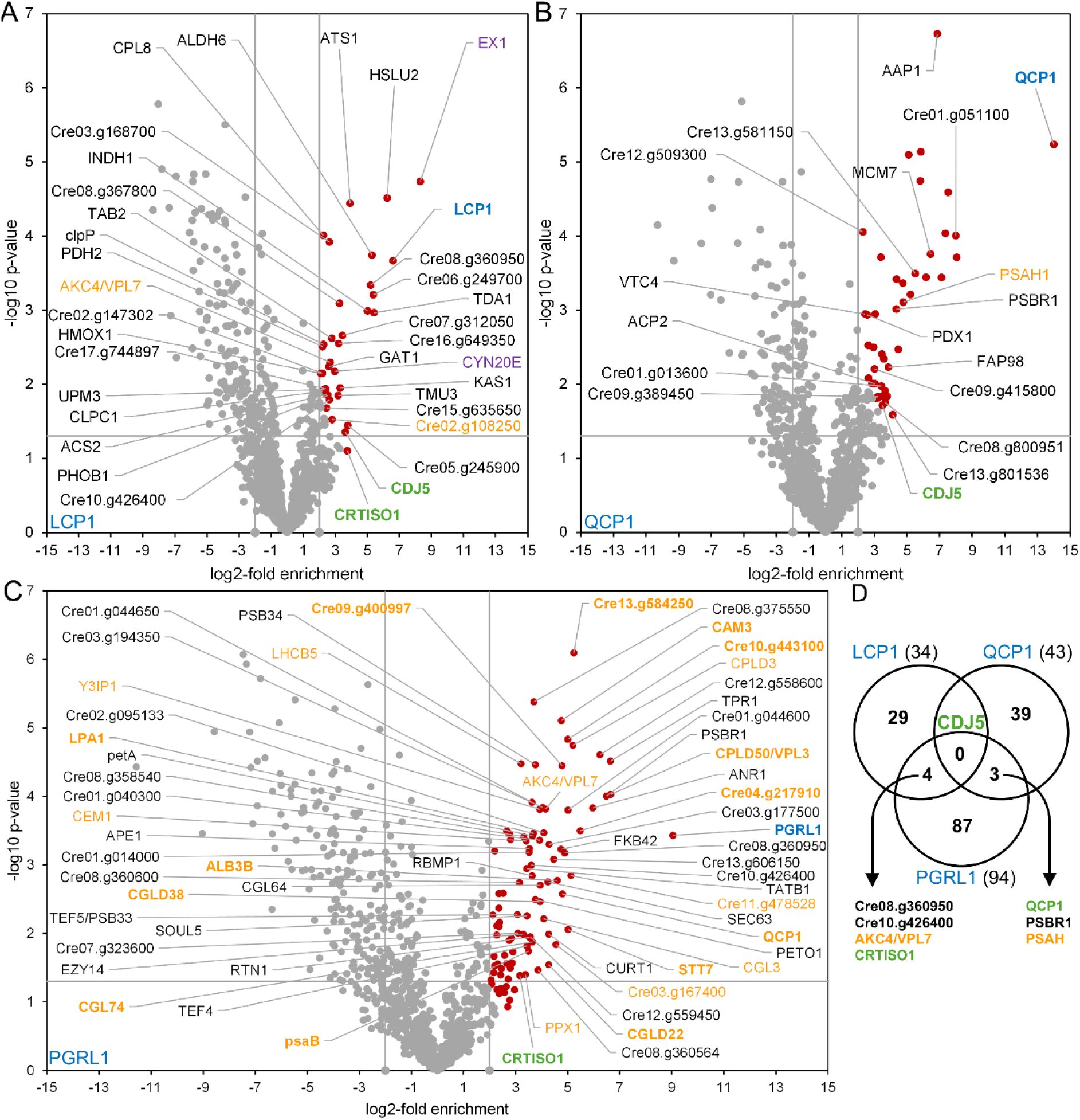
Results of reciprocal proximity labeling for LCP1, QCP1, and PGRL1 found in the CDJ5 proxiomes. (A-C) Volcano plots of the proxiomes of LCP1, QCP1, and PGRL1. Streptavidin bead eluates of the samples shown in Fig. 5C were analyzed by mass spectrometry. Shown is the log2-fold enrichment between proteins in the LCP1_TID_, QCP1_TID_ and PGRL1_TID_ proxiomes versus the mCh_TID_#4 proxiome. Proteins significantly enriched (q ≤ 0.05, enrichment ≥ 4) are shown as red data points. In the QCP1 dataset only proteins are labeled that were predicted to be plastid localized by at least one of six prediction programs to account for the putatively truncated TurboID in the cytoplasm. In the PGRL1 dataset only proteins showing an enrichment ≥ 8 are labeled to reduce complexity. The baits are shown in blue, proteins enriched with all three CDJ5 variants in green, proteins enriched with CDJ5-AAA in orange and those enriched also with CDJ5-WT in bold, and proteins enriched with CDJ5-SSS in purple. (D) Summary of results. The Venn diagram shows the number of proteins enriched with LCP1, QCP1, and PGRL1 and their overlap. The IDs of proteins in the overlaps are given using the same color code as in the volcano plots.

43 proteins were significantly enriched with QCP1_TID_ (excluding QCP1). Its proxiome is characterized by the presence of multiple proteins with proposed localization to the nucleus/cytoplasm and by the extreme enrichment of QCP1 by 16,533-fold (Fig. 6B; Supplemental Dataset S2). We attribute the strong enrichment of nuclear/ cytosolic proteins to the truncation of TurboID from a fraction of the QCP1_TID_ fusion protein (Fig. 5B) which, as assumed for the QCP1_mNG_ fusion, probably has occurred in the cytosol. Since the enrichment is calculated against mCherry_TID_ in the chloroplast, proteins biotinylated by the truncated TurboID in the cytosol will end up as highly enriched. In Fig. 6B we have therefore only labeled proteins that were predicted to localize to the chloroplast by at least one of six prediction programs. The extreme enrichment of the QCP1_TID_ bait is not due to the extreme enrichment of few QCP1 peptides remaining with the truncated cytosolic TurboID, as all peptides identified for QCP1 were highly enriched. Incomplete import of QCP1_TID_ into the chloroplast is also unlikely, since the predicted N-terminal transit peptide was not covered by any peptide while virtually the entire remaining sequence was covered (Supplementary Fig. S6), suggesting that the protein was imported quantitatively. Possibly the strong enrichment of QCP1_TID_ is due to its localization to a chloroplast microcompartment that is barely accessible for mCherry_TID_. CDJ5 was enriched 11.3-fold. In addition to CDJ5, only PSI subunit PSAH (27.3-fold enriched) was also found in the proxiomes of the CDJ5-WT and CDJ5-AAA variants.

Reanalysis of the previously published PGRL1 proxiome (Kreis *et al*., 2023a) with MSFragger resulted in 94 significantly enriched proteins (excluding PGRL1) of which 85% were predicted to be chloroplast-localized by at least one of six prediction programs (Fig. 6C, D; Supplementary Dataset S2). Of the 43 proteins found previously in the PGRL1 proxiome after evaluation with MaxQuant, 33 were also found after reanalysis with MSFragger. The 10 missing proteins were all enriched, but this was below the 4-fold enrichment threshold set here (previously: 2-fold). Among these ten missing proteins was VIPP1 (3-fold enriched, q = 0.052). Like in the MaxQuant evaluation, PGRL1 was 528-fold enriched. CDJ5 was non-significantly enriched only 1.6-fold in the PGRL1_TID_ proxiome which compared to the >11-fold enrichment with LCP1_TID_ and QCP1_TID_ explains why CDJ5 was detected immunologically in the LCP1_TID_ and QCP1_TID_ streptavidin eluate but not in that of PGRL1_TID_ (Fig. 5C).

Two major functional categories were covered in the previous and current PGRL1 proxiome evaluations (Kreis *et al*., 2023a) (Fig. 6C). The first of which is regulation of photosynthetic electron flow with ANR1, PETO, FNR1, psaA, LHCA4, LHCB5, and LHCBM5 which have been found in complex with PGRL1 (Iwai *et al*., 2010; Takahashi *et al*., 2016; Terashima *et al*., 2012) as well as STT7 and PBCP1 involved in state transitions. The new evaluation added further components of the PGRL1-containing CEF supercomplex (Iwai *et al*., 2010) including PSI subunits psaB, PSAD, PSAH, and PSAK as well as Cyt b_6_/f subunit petA.

The second major functional category in the PGRL1 proxiome is thylakoid membrane protein complex biogenesis with CGLD22 and possibly Cre13.g584250 (ATP synthase), Y3IP1 (PSI) (Albus *et al*., 2010), TEF5 (PSII) (Lang *et al*., 2025) and LPA1 (PSII). The new evaluation added RBD1 and PSB34 (PSII) (Zabret *et al*., 2021) as well as ALB3B, TATB and SecY1 (membrane insertases/translocases) (Schuenemann *et al*., 1999; Zinecker *et al*., 2020). The new evaluation also revealed further functional groups with at least three proteins: these are carotenoid biosynthesis with P450 protein CYP97C3 and CRTISO adding to previously identified P450 protein CYP97B6 (Lohr, 2023); and chlorophyll biogenesis adding CHLM1 and CGL74 to previously identified proteins PPX1 and CHLP1 (Willows *et al*., 2023). Overall, there was only little overlap between the proxiomes of LCP1, QCP1 and PGRL1 (Fig. 6D). CDJ5 was the only protein commonly found in the proxiomes of LCP1 and QCP1. The proxiomes of LCP1 and PGRL1 shared only four proteins including CRTISO1 and AKC4/VPL7 identified previously in proximity of VIPP1/2 (Kreis *et al*., 2023a) as well as two proteins of unknown function. The QCP1 and PGRL1 proxiomes shared only three proteins, which are QCP1 and photosystem subunits PSBR1 and PSAH.

### The PGRL1 proxiome overlaps largely with the proxiomes of CDJ5 variants capable of binding a 4Fe-4S cluster

Of the 94 proteins in the PGRL1 proxiome, 34 were also found in the largely overlapping proxiomes of CDJ5-WT and CDJ5-AAA (Fig. 6C; Fig. 7). Among 12 proteins of unknown function, these shared proteins belong mostly to the two main functional categories identified in the PGRL1 proxiome: regulation of photosynthetic electron flow with members of the PGRL1-containing CEF complex (psaA, psaB, PSAH, LHCB5) and regulators of state transitions (STT7, PBCP1); and thylakoid membrane protein biogenesis (LPA1, RBD1, Y3IPI, ALB3B, CGLD22 and possibly Cre13.g584250) as well as pigment biosynthesis (CRTISO1, CYP97B6, CGL74, PPX1).

**Fig. 7.**
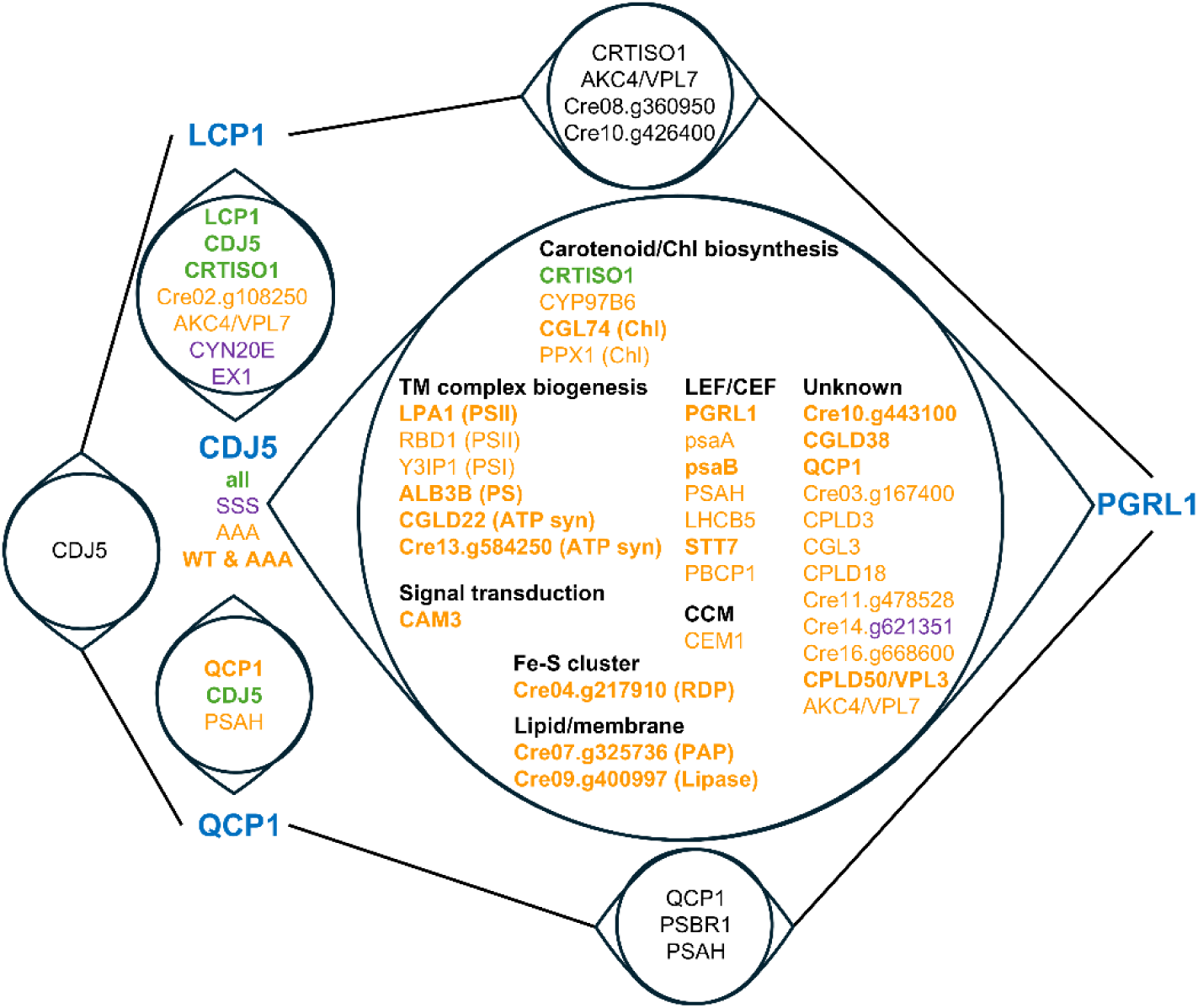
Overlap of proxiomes of CDJ5 variants, LCP1, QCP1, and PGRL1. Only proteins identified in the proxiomes of the three CDJ5 variants are colored as follows: proteins in the proxiomes of all three variants are shown in green; proteins in the proxiome of CDJ5-AAA in orange and those present in the proxiome of CDJ5-WT as well in bold; proteins enriched in the proxiomes of CDJ5-SSS in purple. Proteins present in the CDJ5-WT/AAA and PGRL1 proxiomes are sorted according to common functional categories (black).

## Discussion

In this work, we employed TurboID-mediated proximity labeling to obtain insights into the function of CDJ5, since all our attempts to obtain *cdj5* knock-out or knock-down lines via CRISPR/Cas9 or amiRNA failed. We found HSP70B in the proxiome of WT CDJ5 and in the proxiomes of CDJ5 variants impaired in stimulating the ATPase activity of HSP70B (CDJ5- AAA) or incapable of binding a 4Fe-4S cluster (CDJ5-SSS) (Figs. 2D and 3). This finding adds CDJ5 as another bona-fide cochaperone of stromal HSP70B in addition to CDJ1-4, which have been shown to interact with HSP70B previously (Dorn *et al*., 2010; Liu *et al*., 2007; Liu *et al*., 2005; Veyel *et al*., 2014; Willmund *et al*., 2008). HSP90C was also found to be significantly enriched in the proxiomes of all three CDJ5 variants, corroborating its cooperation with HSP70B and J-domain cochaperones (Schroda and Mühlhaus, 2009; Willmund *et al*., 2008).

We found that the proxiomes of CDJ5 variants capable of binding a 4Fe-4S cluster (CDJ5-WT and CDJ5-AAA) overlap almost entirely but differ largely from the proxiome of CDJ5-SSS incapable of binding a 4Fe-4S cluster (Fig. 3). This finding suggests that a functional 4Fe-4S cluster (or its redox state?) determines the localization of CDJ5 to a specific chloroplast microcompartment. The largest functional groups in the proxiomes of CDJ5 variants capable of binding a 4Fe-4S cluster consisted of proteins involved in the biogenesis of thylakoid membrane protein complexes (photosystems, Cyt b_6_/f complex, ATP synthase) and their pigments (carotenoids and chlorophyll) as well as in the regulation of photosynthetic electron transport (CEF, state transitions in LEF) (Fig. 3). The proxiome of CEF component PGRL1 largely overlapped with that of CDJ5-WT and CDJ5-AAA but not with that of CDJ5- SSS (Figs. 6 and 7), suggesting that CDJ5 is present in the same chloroplast microcompartment as PGRL1 and that this depends on its capability to bind a 4Fe-4S cluster or on the redox state of this cluster. Accordingly, CDJ5-WT_mNG_ and PGRL1_mNG_ share a similar general localization pattern in the chloroplast (Figs. 1E and 4D). These data suggest that CDJ5 with HSP70B/HSP90C might play a role in the regulation of thylakoid membrane protein complex biogenesis and photosynthetic electron transport via its redox-active 4Fe-4S cluster. The proxiomes of LCP1 and QCP1 also found in the proxiomes of the CDJ5 variants differed substantially from the PGRL1 proxiome, with only few shared proteins (Figs. 6 and 7). This finding indicates that CDJ5 is present in several different chloroplast microcompartments with distinct protein environments. LCP1 consists only of the N domain of LonA proteases which are involved in the selection of damaged and some short-lived regulatory proteins for their subsequent degradation (Wlodawer *et al*., 2022). Since LCP1 lacks the central ATPase domain and the C-terminal proteolytic domain, it is unclear what purpose LCP1 with the N domain alone could serve. It is interesting that we found class II and class I HSP100 members HSLU2 and ClpC1, respectively, as well as subunit clpP of the proteolytic chamber in the proxiome of LCP1 (Fig. 6A). Here it is tempting to speculate that substrates recognized by LCP1 could be delivered to these HSP100 members coupled with the clpP proteolytic chamber for degradation. Although HSLU2 is supposed to be localized to mitochondria (Schroda and deVitry, 2023), programs Plant-mSubP and BacCelLo predicted a chloroplast localization. Striking is the 315-fold enrichment of the EXECUTER1 (EX1) homolog in the LCP1 proxiome, which was also enriched 68.6-fold in the CDJ5-SSS proxiome. This suggests a role of LCP1 and a specific redox state of the 4Fe-4S cluster mimicked in CDJ5-SSS in ROS signaling. It was proposed that EX1-associated proteins may release signaling molecules upon EX1 proteolysis following oxidation of a specific Trp residue in its DUF3506 ^1^O_2_ sensor domain (Kim, 2020). While an interaction of LCP1 and EX1 suggests a role of LCP1 in ^1^O_2_ signaling in *Chlamydomonas*, Arabidopsis LCP1 was shown to interact and inhibit Trx-y2 that activates peroxidase scavenging of H_2_O_2_, resulting in elevated H_2_O_2_ concentrations (Shin *et al*., 2020). QCP1, which contains a domain that is structurally very similar to the PSBQ protein of PSII, has PSAH in its proxiome which is present also in the proxiomes of CDJ5 and PGRL1, but not in that of LCP1 (Fig. 7). For yet unknown reasons, PSI subunit PSAH was found to be particularly enriched in pyrenoid tubules (Mackinder *et al*., 2017). Accordingly, in addition to co-localizing with chlorophyll autofluorescence, QCP1_mNG_ fluorescence was found in the pyrenoid in structures that could represent pyrenoid tubules, like fluorescence from mNG fused to CDJ5, LCP1, and PGRL1 (Figs. 1E, 4D; Supplementary Figs. S1, S7, S8).

### Limitations and strengths of the proximity labeling approach

PGRL1, LCP1, and QCP1 were enriched in the CDJ5-WT proxiome by 9.2-, 25-, and 7.5-fold, respectively, while CDJ5-WT was reciprocally enriched only in the LCP1 and QCP1 proxiomes by 12.4- and 11.3-fold, respectively (Figs. 3, 5C, 6). In the PGRL1 proxiome, CDJ5 was hardly enriched (1.6-fold, not significant), however, the proxiome of PGRL1 overlapped largely with those of CDJ5-WT and CDJ5-AAA, indicating that functional CDJ5 and PGRL1 do act in the same chloroplast microcompartment. How can this discrepancy be explained? One possible explanation is that for sterical reasons the cloud of activated biotin generated by CDJ5_TID_ reaches PGRL1 but not vice versa. Alternatively, since the enrichment is always calculated against biotinylation by the mCherry_TID_ control, a closer proximity of mCherry_TID_ to CDJ5 than to PGRL1 would result in a stronger non-specific biotinylation of CDJ5 and thus reduce the contrast. Both scenarios could be related to the fact that PGRL1 is a membrane-bound protein with two transmembrane helices, whereas CDJ5, LCP1, QCP1, and mCherry are not membrane-bound. Here, the overlapping proxiomes might be better indicators of proximity than reciprocal enrichment of the two target proteins.

Another limitation of our proximity labeling approach became evident with QCP1_TID_, where TurboID was obviously cleaved off from a fraction of QCP1_TID_ before import into the chloroplast (Fig. 5B, C). This apparently occurred also for the QCP1_mNG_ fusion resulting in mNG fluorescence originating also from the cytosol (Fig. 4C; Supplementary Fig. S9). This resulted in a strong enrichment of many cytosolic proteins since the enrichment is calculated against the mCherry_TID_ control targeted quantitatively to the chloroplast. The chloroplast prediction of 85-100% of the proteins in the proxiomes of the CDJ5 variants, LCP1, and PGRL1 by at least one of six prediction programs suggests that there was no cleavage of the TurboID tag prior to chloroplast import for these fusion proteins. The proteins not predicted to localize in the chloroplast could have incorrect annotations of their N-termini resulting in erroneous transit peptide sequences. This includes alternative splicing events and alternatively used start codons. Alternative import pathways could also play a role (Villarejo *et al*., 2005). For sure, the dual targeting of proteins to several compartments was recently revealed as a frequent phenomenon in *Chlamydomonas* (Wang *et al*., 2023). In any case, the wealth of information gained by proximity labeling on the localization of proteins to specific microcompartments is definitely a strength of the proximity labeling approach and will help shaping the protein localization atlas of *Chlamydomonas*. For example, the most enriched protein in the proxiomes of all three CDJ5 variants was CCP1, which by fractionation experiments was reported to be localized to chloroplast envelopes (Ramazanov *et al*., 1993) but previously via heterologous expression with a C-terminal mVenus tag shown to localize in mitochondria (Mackinder *et al*., 2017). Our detection of CCP1 in the proxiome of CDJ5 points to the presence of at least a fraction of CCP1 in the chloroplast.

## Limitations of our study

Since we were not able to obtain *cdj5* knock-out or knock-down lines, we could not verify the functionality of CDJ5 with C-terminal 3xHA, mNG, or TurboID tags. The functionality of C-terminal fusion proteins can only be implied by the specific enrichment of the expected chaperone partners HSP70B (and HSP90C) in the proxiomes of CDJ5_TID_. Since CDJ5 has not been detected in recent proteomics analyses, it appears to be weakly expressed (Schroda and deVitry, 2023). Therefore, it is not clear whether the localization and thus the proxiome of the overexpressed CDJ5 fusion proteins is the same as for the native protein. This is however suggested by the distinct proxiomes of CDJ5-WT and CDJ5-AAA versus CDJ5-SSS with destroyed binding sites for the 4Fe-4S cluster. Although these limitations apply to any study involving the heterologous overexpression of target proteins, they should be kept in mind.

## Supplementary data

The following supplementary data are available at JXB online.

**Table S1.** Primers used for cloning.

**Fig. S1.** Localization of CDJ5-WT_mNG_ in transformants #3 and #4 generated with pMBS1209.

**Fig. S2.** Analysis of transformants producing mNG in the chloroplast stroma.

**Fig. S3.** Analysis of transformants producing mNG in the cytosol.

**Fig. S4.** Analysis of cpUPR marker accumulation in transgenic lines producing 3xHA and TurboID fusion proteins.

**Fig. S5.** Amino acid sequence and structural alignments for LCP1.

**Fig. S6.** Amino acid sequence and structural alignments for QCP1.

**Fig. S7.** Localization of PGRL1_mNG_ in transformants #7 and #9 generated with pMBS1239. **Fig. S8.** Localization of LCP1_mNG_ in transformants #9 and #11 generated with pMBS1241.

**Fig. S9.** Localization of QCP1_mNG_ in transformant #7 generated with pMBS1240.

**Dataset S1.** Proximity labeling results of CDJ5-WT, CDJ5-AAA, and CDJ5-SSS.

**Dataset S2.** Proximity labeling results of LCP1, QCP1, and PGRL1.

## Acknowledgements

We would like to thank Lara Peters for her help in the initial stages of the project.

## Author contributions

K.K. and M.M. performed all experiments supported by J.N. S.Z. generated the constructs for cytosolic and chloroplast mNeonGreen controls. S.G. implemented the localization algorithms.

F.S. performed the mass spectrometry analyses. M.S. conceived and supervised the project and wrote the article with contributions from all authors.

## Data availability statement

All data supporting the findings of this study are available within the paper and within its supplementary materials published online.

